# Cellpose as a reliable method for single-cell segmentation of autofluorescence microscopy images

**DOI:** 10.1101/2024.06.07.597994

**Authors:** Jeremiah M Riendeau, Amani A Gillette, Emmanuel Contreras Guzman, Mario Costa Cruz, Aleksander Kralovec, Shirsa Udgata, Alexa Schmitz, Dustin A Deming, Beth A Cimini, Melissa C Skala

## Abstract

Autofluorescence microscopy uses intrinsic sources of molecular contrast to provide cellular-level information without extrinsic labels. However, traditional cell segmentation tools are often optimized for high signal-to-noise ratio (SNR) images, such as fluorescently labeled cells, and unsurprisingly perform poorly on low SNR autofluorescence images. Therefore, new cell segmentation tools are needed for autofluorescence microscopy. Cellpose is a deep learning network that is generalizable across diverse cell microscopy images and automatically segments single cells to improve throughput and reduce inter-human biases. This study aims to validate Cellpose for autofluorescence imaging, specifically from multiphoton intensity images of NAD(P)H. Manually segmented nuclear masks of NAD(P)H images were used to train new Cellpose models. These models were applied to PANC-1 cells treated with metabolic inhibitors and patient-derived cancer organoids (across 9 patients) treated with chemotherapies. These datasets include co-registered fluorescence lifetime imaging microscopy (FLIM) of NAD(P)H and FAD, so fluorescence decay parameters and the optical redox ratio (ORR) were compared between masks generated by the new Cellpose model and manual segmentation. The Dice score between repeated manually segmented masks was significantly lower than that of repeated Cellpose masks (p<0.0001) indicating greater reproducibility between Cellpose masks. There was also a high correlation (R^2^>0.9) between Cellpose and manually segmented masks for the ORR, mean NAD(P)H lifetime, and mean FAD lifetime across 2D and 3D cell culture treatment conditions. Masks generated from Cellpose and manual segmentation also maintain similar means, variances, and effect sizes between treatments for the ORR and FLIM parameters. Overall, Cellpose provides a fast, reliable, reproducible, and accurate method to segment single cells in autofluorescence microscopy images such that functional changes in cells are accurately captured in both 2D and 3D culture.

## Introduction

Autofluorescence provides an innate source of molecular contrast that is attractive for imaging cells within intact, unaltered systems. Intrinsic sources of contrast such as autofluorescence also avoid the effects of fluorescent labels and genetic modifications, which may interfere with cell function^1,2^. Therefore, autofluorescence microscopy continues to be used to study *in vivo* and *in vitro* models across numerous applications including cancer research, stem cell biology, and metabolism research^1,3^. Autofluorescence signals are generally brightest in the cell cytoplasm, which provides an opportunity to perform single cell segmentation and investigate cellular heterogeneity. However, autofluorescence microscopy suffers from relatively low signal-to-noise ratios (SNR) that make cell segmentation challenging.

Software such as CellProfiler, Fiji/ImageJ, and BioImageXD have been developed for cell segmentation in microscopy, though many such algorithms have been optimized for images that use extrinsic labels with high SNR^4–7^. However, in cases where SNR is low such as in autofluorescence microscopy, segmentation methods including watershed and thresholding can be unreliable. Therefore, a new set of segmentation tools are needed for low SNR applications.

Recent advances in deep learning-based segmentation models have increased the speed, accuracy, and efficiency of segmentation while producing increasingly generalizable models with limited need for supervision. Cellpose is one such deep learning-based cell segmentation algorithm that uses training sets of manually segmented images to learn segmentation patterns for new, unseen images^8^. This automated method minimizes personnel time and human-to-human variances in cell segmentation, thereby increasing the throughput of image analysis. As autofluorescence microscopy becomes more widely adopted, it is important to develop and validate automated methods, such as Cellpose, to mitigate human biases, improve agreement and reproducibility, and increase the throughput of single cell analysis.

Given that convolutional neural networks are traditionally trained on images with clear staining, there is need to validate these approaches on more challenging data sets such as those acquired from autofluorescence microscopy. To achieve this, we trained a Cellpose model on a selection of manually segmented nuclei from autofluorescence intensity images of the metabolic co-enzymes NADH and NADPH, collectively referred to as NAD(P)H due to their overlapping spectral properties^9^. We then examined the reproducibility of the Cellpose model compared to manual segmentation on new sets of images. We further compared changes in functional autofluorescence variables for manual and Cellpose segmented images, including fluorescence lifetime imaging microscopy (FLIM) of NAD(P)H and flavin adenine dinucleotide (FAD), another autofluorescent metabolic co-enzyme. Both NAD(P)H and FAD have known distinct lifetimes in the free and protein-bound states, therefore FLIM provides insight into protein-binding activities for these molecules, and as a result, the metabolic activity of the cells^9,10^. Additionally, we assessed the optical redox ratio (fluorescence intensities of NAD(P)H / [NAD(P)H + FAD]), which reflects changes in redox status within cells^11–13^. Here, FLIM is performed with two-photon (2P) microscopy to enable optical sectioning at deeper imaging depths in 3D samples compared to single-photon excitation^14^. We sought to investigate the accuracy of Cellpose segmentation performance on NAD(P)H intensity images in both two-dimensional (2D) cell lines and three-dimensional (3D) patient-derived cancer organoids (PDCOs) with varying morphologies. Through this study, we aim to provide a deeper understanding of the strengths and limitations of Cellpose for single-cell segmentation and how this method compares to manual segmentation of autofluorescence microscopy images.

## Results

### Similarity of Cellpose and manual masks

Representative NAD(P)H intensity images of 2D PANC-1 cells overlain with their respective masks indicate strong consensus between manual and Cellpose masks **(Fig. 1a)**. To measure differences between manual and Cellpose masks, the Dice similarity coefficient (DSC), or Dice score was used. This metric measures the amount of overlap between two sets of masks by multiplying the area of overlap of the two images by two, then dividing by the total area of both images **(Fig. 1b)**, thus penalizing false positives. When the manually produced masks (M1) are compared with the Cellpose segmented masks (C1), an average Dice score of 0.74 is found for the M1-C1 comparison **(Fig. 1b)**. To further categorize the reproducibility of masks, images were rotated by 90 degrees and segmented again (M2 and C2) resulting in an average Dice score of 0.78 for the manually segmented initial orientation and rotated images, M1-M2 **(Fig. 1c)**. This is in agreement with prior work, where masks produced by different humans achieved a two-observer maximal Dice score of 0.76 on a 0-1 scale^8^. Importantly, Cellpose achieved a higher average Dice score of 0.87 for the automatic segmentation of initial orientation and rotated images, C1-C2 **(Fig. 1c)**, indicating greater reproducibility between Cellpose masks compared to manual segmentation twice by the same observer.

**Figure 1.**
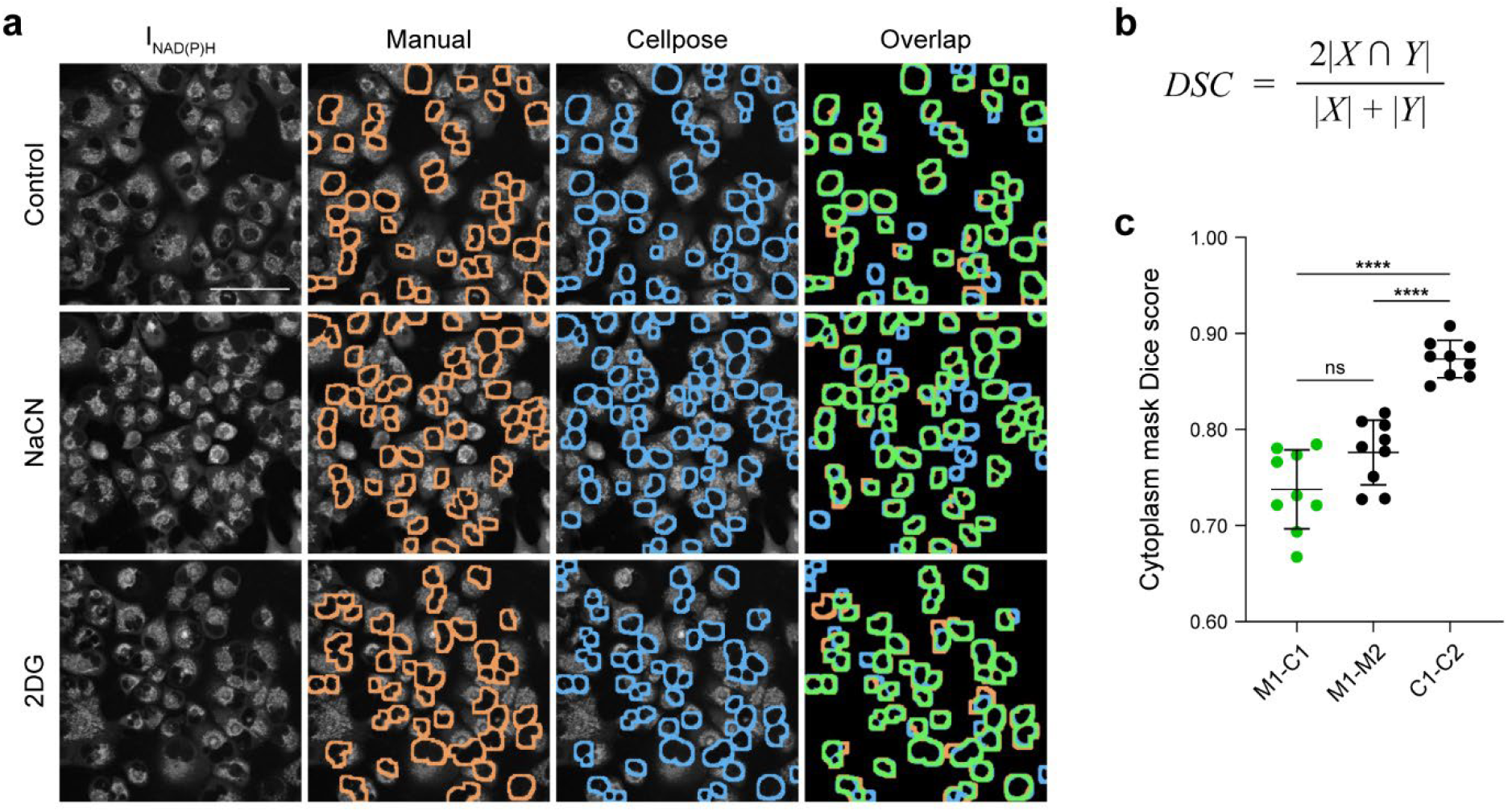
Cellpose masks agree with manual masks while achieving greater reproducibility in 2D culture. (a) Representative two-photon microscopy images of NAD(P)H intensity (left) of PANC-1 cells with or without treatment with inhibitors of oxidative phosphorylation (NaCN 4mM; 30 min) or glycolysis (2DG 10mM; 120 min). Masks are overlain for manual segmentation (orange) and Cellpose segmentation (blue) where more overlap (green) results in a higher dice score. Cytoplasm area is defined as 8 µm propagated from the nuclear mask. Scale bar = 100 µm. (b) Dice Similarity Coefficient (DSC), or Dice Score, is calculated by multiplying the green pixel area by two, then dividing that value by the sum of the pixel areas of the blue and orange regions. (c) Dice scores compare Manual [M] and Cellpose [C] generated masks where M2 and C2 images have been rotated 90° with respect to M1 and C1 images such that M1-M2 compares manual reproducibility and C1-C2 compares Cellpose reproducibility. Each dot represents one image. Center line is mean; error bars are standard deviation. Control n = 3 images, 146 nuclei (manual), 160 nuclei (Cellpose); NaCN n = 3 images, 143 nuclei (manual), 217 nuclei (Cellpose); 2DG n = 3 images, 143 nuclei (manual), 173 nuclei (Cellpose). ****P<0.0001, ns = P>0.05, Ordinary one-way ANOVA.

While 2D cells are relatively homogenous in shape and size across treatment conditions, four distinct morphologies were found in the 3D PDCOs **(Fig. 2, Supp. Table 1)**. These morphologies were characterized by their shape, appearing either circular or irregular, and their contents, either densely filled with cells or void of cells. Variability in PDCO morphologies are well documented and can arise through a variety of mechanisms ranging from the genetic diversity in the sample to the randomness of the self-organization process as well as their developmental stage^15–17^. These morphological variances can be clearly seen in 2P NAD(P)H intensity images **(Fig. 2a)**. Combining all PDCO morphologies demonstrates similar reproducibility for manual vs Cellpose segmentation (M1-C1 Dice score = 0.68) and manual vs manual segmentation (M1-M2 Dice score = 0.66), with greater reproducibility between Cellpose masks (C1-C2 Dice score = 0.92) **(Fig. 2b)**. Therefore, in these 3D PDCOs, Cellpose masks are more reproducible than manual masks, which is consistent with prior studies and the findings in 2D cells^18^ **(Fig. 1)**.

**Figure 2.**
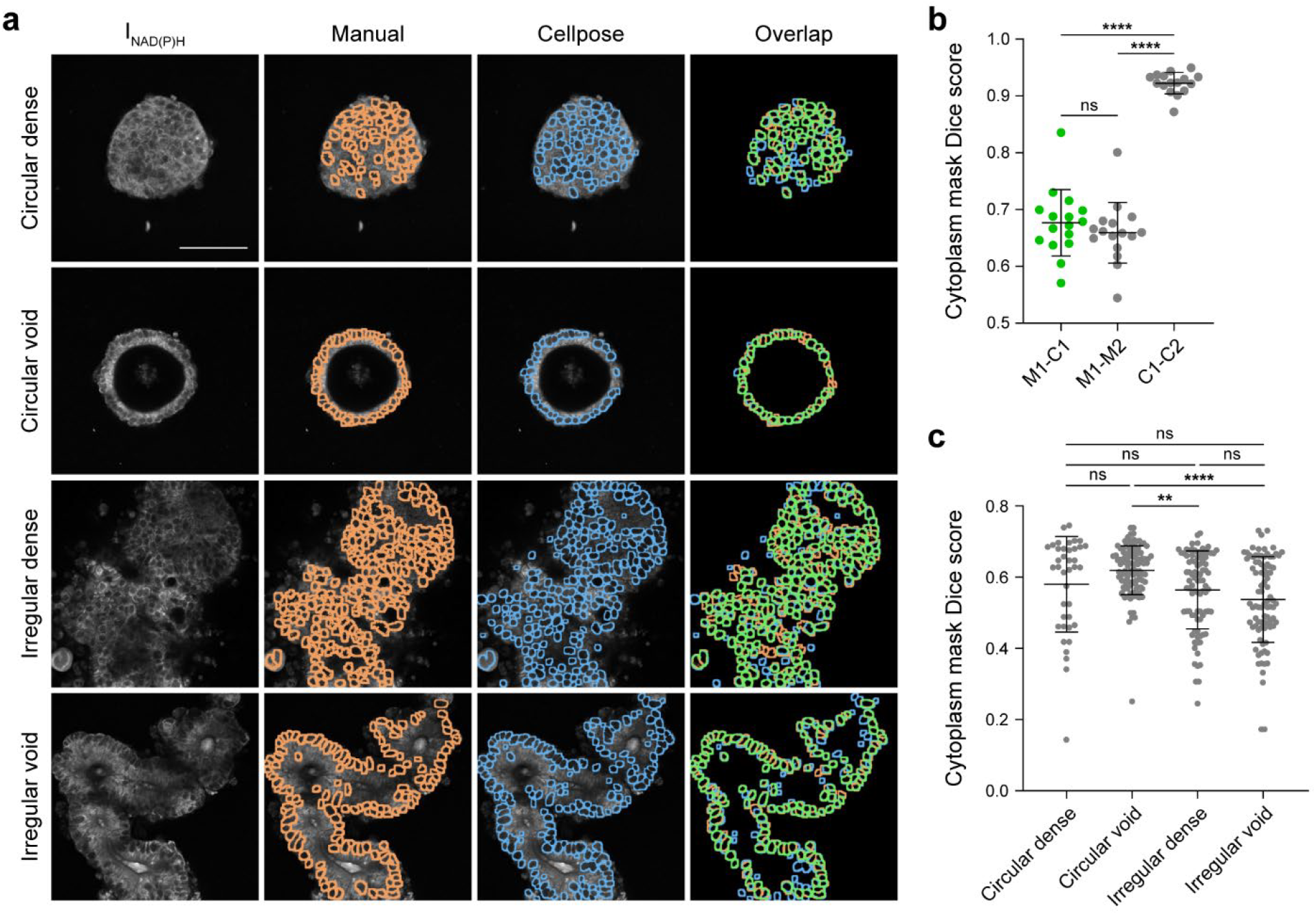
Cellpose masks agree with manual masks while achieving greater reproducibility in optical cross sections of 3D PDCOs. (a) Representative NAD(P)H intensity images of rectal PDCOs (images acquired from Kratz et al.^27^) with morphologies defined as circular or irregular and dense or void. Masks are overlain for manual segmentation (orange) and Cellpose segmentation (blue) where more overlap (green) results in a higher dice score. Cytoplasm area is defined as 8 µm propagated from the nuclear mask. Scale bar = 100 µm. (b) n=16 organoids representative of the irregular dense morphology, chosen for their complexity. Dice scores compare manual [M] and Cellpose [C] segmented masks where M2 and C2 images have been rotated 90°. Cellpose segmented masks display more similarity when the image is rotated and segmented again [C1-C2] than when a human performs the same task [M1-M2]. (c) Comparison of Cellpose segmentation and manual segmentation of PDCOs across PDCO morphologies. The two-observer maxima for the Dice score measurement is 0.76 (see M1-C1 and M1-M2), similar to inter-observer Dice scores [1]. n = 28 circular dense images, 2973 nuclei (manual), 2544 nuclei (Cellpose); 81 circular void images, 7868 nuclei (manual), 7294 nuclei (Cellpose); 70 irregular dense images, 9318 nuclei (manual), 9048 nuclei (Cellpose); and 65 irregular void images, 7277 nuclei (manual), 8334 nuclei (Cellpose). Center line is mean and error bars are standard deviation. ****P<0.0001, ns = P>0.05, Ordinary one-way ANOVA.

All PDCO images were acquired from a previously published dataset in Kratz et al.^27^ Due to the varying morphologies and spatial arrangements of the PDCOs, the performance of Cellpose between different morphologies was explored **(Fig. 2c)**. Dice scores comparing the manual vs Cellpose masks (M1-C1) found that three of the morphologies show similar agreement (Dice score = 0.54-0.58) while the circular void morphology, which contains fewer cells on average, has a slightly higher agreement (Dice score = 0.62). These varying PDCO morphologies can be found in many cancer types including breast and pancreatic cancers, providing a robust and varied testing set for the generalizability of Cellpose across different shapes, sizes, and cell environments^19,20^.

In addition to providing similar segmentation as manual segmentation, Cellpose segments images several orders of magnitude faster **(Supp. Fig. 1)**. While our expert will spend roughly 11 minutes to segment 100 nuclei, Cellpose can perform the same number of nuclei segmentations in less than a second **(Supp. Fig. 1)**. It is worth noting that these results were only compiled for one human and segmentation time will vary from person to person, while automation time depends on hardware and software systems. However, these results clearly indicate that the speed and reproducibility of automated Cellpose segmentation is unachievable by humans as demonstrated in numerous prior studies^8,21,22^.

### Metabolic parameters are preserved using Cellpose segmentation

Differences between Cellpose and manual segmentation were investigated with treatment in 2D and 3D cultures using autofluorescence FLIM parameters as treatment response measurements, where a decrease in ORR and NAD(P)H mean lifetime (τ_m_) indicates treatment response^23^. Autofluorescence FLIM parameters include the optical redox ratio (ORR), NAD(P)H τ_m_, and FAD τ_m_. PANC-1 cells were treated with either 4 mM sodium cyanide (NaCN) for 30 minutes or 10 mM 2-deoxy-glucose (2DG) for 2 hours. These two metabolic inhibitors have known effects on autofluorescence FLIM parameters which we replicate in our 2D data^24–26^. Comparison between the values produced from the masks created by Cellpose and those created manually **(Fig. 3a-c)** indicates strong agreement in autofluorescence FLIM parameter changes. Outputs from Cellpose and manual segmentation are similar with an R^2^ value of 0.984 for NAD(P)H τ_m_, 0.989 for FAD τ_m_, and 0.964 for ORR. Aside from maintaining near-identical values, Cellpose also retains similar single-cell distributions **(Fig. 3d-f)** while maintaining the same effect sizes between treatment groups. This suggests that in 2D cultures, Cellpose performs exceptionally well.

**Figure 3.**
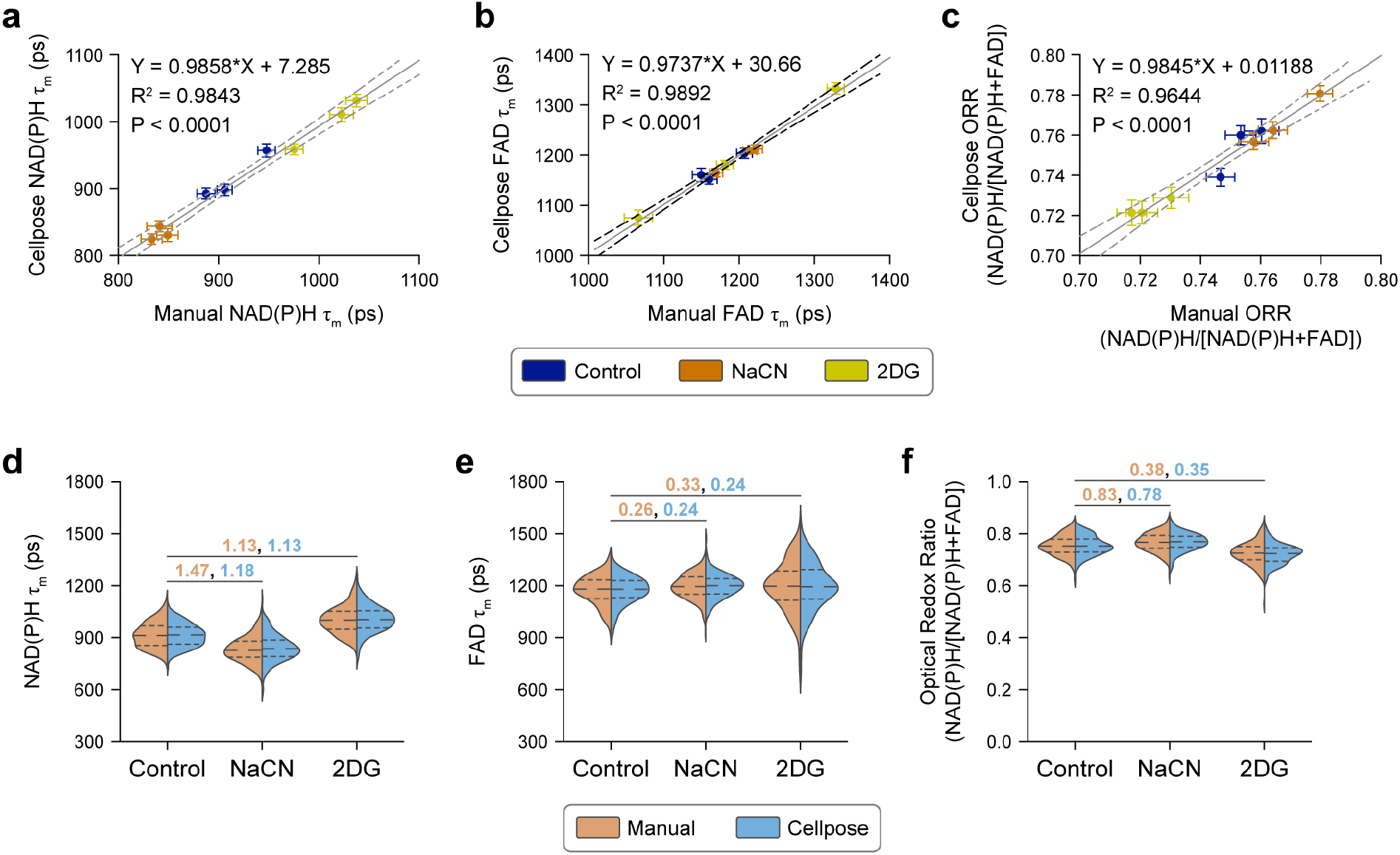
Autofluorescence FLIM parameters from Cellpose and manual segmentation are highly correlated for 2D cultured PANC-1 cells under 3 treatment conditions. PANC-1 cells were left untreated or treated with an inhibitor of oxidative phosphorylation (NaCN 4 mM; 30 min) or glycolysis (2DG 10 mM; 120 min). (A-C) Autofluorescence FLIM parameters generated from manual segmented images and Cellpose segmented images are highly correlated (R^2^>0.9643). Each dot represents one image, colored by treatment condition, where error bars represent standard error of the mean. Solid black diagonal line is linear regression and dashed lines represent 95% confidence bands. P value determined by ratio T test of Spearman correlation values. (D-F) Glass’s Delta effect sizes between control and treatment groups are maintained with both Cellpose segmentation and manual segmentation. Cellpose segmentation retains cell distributions observed with manual segmentation. Dashed lines separate quartiles. Control n = 3 images, 146 nuclei (manual), 160 nuclei (Cellpose); NaCN n = 3 images, 143 nuclei (manual), 217 nuclei (Cellpose); 2DG n = 3 images, 143 nuclei (manual), 173 nuclei (Cellpose). Orange numbers are effect size between manually segmented cells and blue numbers are effect size between Cellpose segmented cells. No statistical difference is measured between the manual and Cellpose segmented groups for the same condition (side-by-side orange and blue violin plots).

Autofluorescence FLIM is also a useful tool for measuring treatment response in 3D PDCO cultures, where cell borders become blurred, and cells become more irregular as they compact into dense PDCO structures^24,27^. Autofluorescence FLIM parameters across multiple treatments remain similar for Cellpose and manual segmentation of PDCOs **(Fig. 4a-c)** with an R^2^ value of 0.98 for NAD(P)H τ_m_, 0.98 for FAD τ_m_, and 0.86 for ORR and appear to maintain similar agreement despite morphological differences **(Supp. Fig. 2)**. Autofluorescence FLIM parameters are also plotted by treatment condition in PDCOs **(Fig. 4d-i)**, which indicates that Cellpose retains similar single-cell distributions to manually segmented images. Responses for individual patients with respect to treatment condition can be found in supplemental materials **(Supp. Fig. 3)**. Autofluorescence FLIM parameters for PDCOs segmented with Cellpose and manual segmentation also maintain similar levels of effect size between treatment groups despite differences between the number of nuclei segmented manually or by Cellpose, demonstrating a high level of performance even in 3D PDCOs **(Fig. 4d-i, Supp. Table. 1)**.

**Figure 4.**
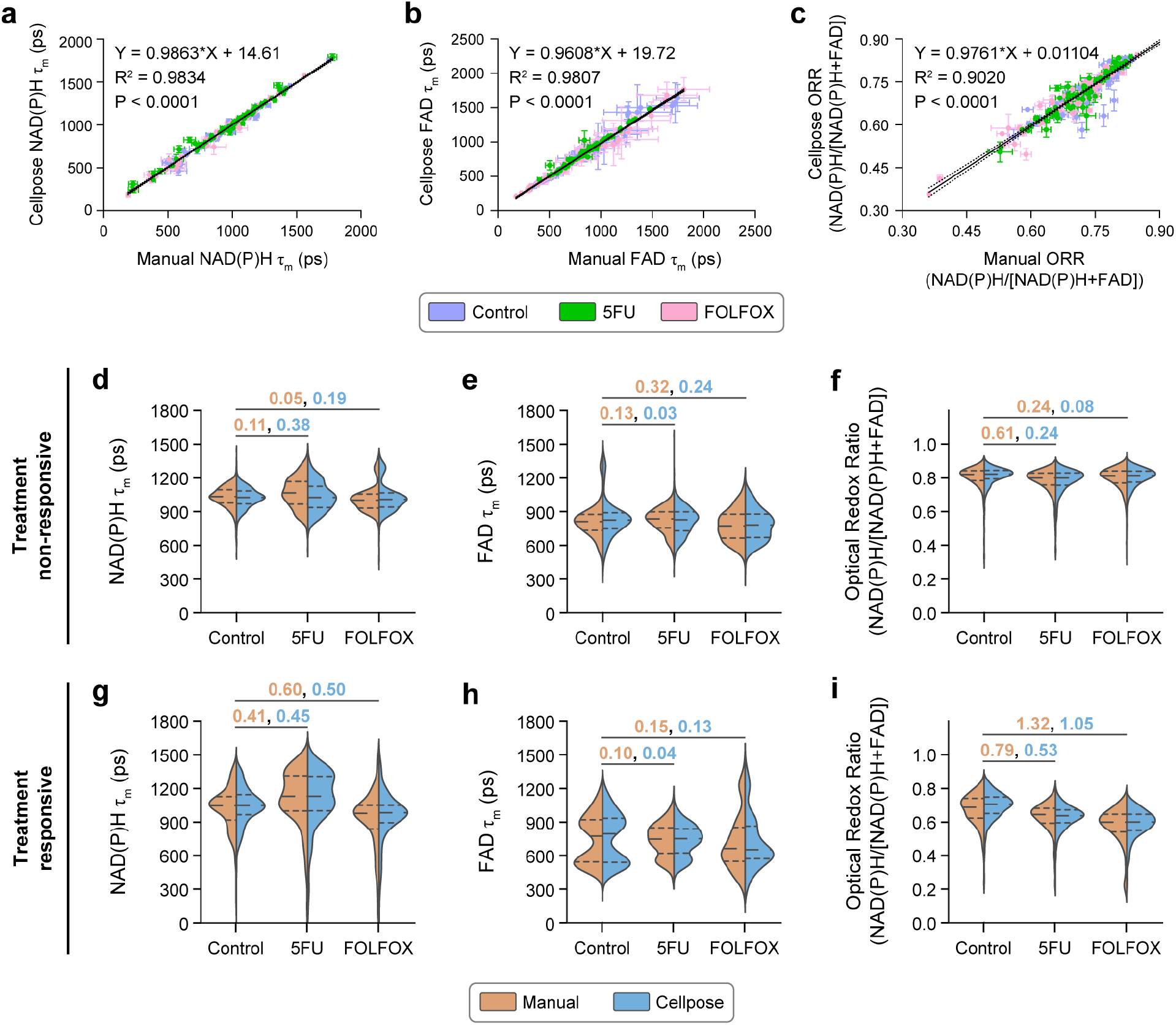
Autofluorescence FLIM parameters from Cellpose and manual segmentation are highly correlated for all 3D colorectal PDCOs across 4 treatment conditions. PDCOs were left untreated or treated with 10 µmol/L of 5FU, or FOLFOX (10 µmol/L of 5FU and 40 mmol/L of oxaliplatin) for 4 days (images acquired from Kratz et al.^27^). (A-C) Autofluorescence FLIM parameters generated from manually segmented images and Cellpose segmented images are highly correlated (R>0.9019). Each dot represents one organoid image, where error bars represent standard error of the mean. Black diagonal line is linear regression and dashed lines represent 95% confidence bands. P value determined by ratio T test of Spearman correlation values. (D-H) Significant differences between control and treatment groups are maintained with both Cellpose segmentation and manual segmentation of (D-F) PDCOs that do not respond to treatment or (G-I) PDCOs that are treatment responsive. Cellpose segmentation retains cell distributions observed within manual segmentation, where dashed lines separate quartiles. Nonresponsive PDCOs (patient 6) Control n = 8 images, 797 nuclei (manual), 979 nuclei (Cellpose); 5FU n = 8 images, 983 nuclei (manual), 1009 nuclei (Cellpose); FOLFOX n = 7 images, 730 nuclei (manual), 816 nuclei (Cellpose). Responsive PDCOs (patient 3) Control n = 9 images, 672 nuclei (manual), 1161 nuclei (Cellpose); 5FU n = 9 images, 635 nuclei (manual), 1042 nuclei (Cellpose); FOLFOX n = 9 images, 611 nuclei (manual), 808 nuclei (Cellpose). Orange numbers are effect size between manually segmented cells and blue numbers are effect size between Cellpose segmented cells. No statistical difference is measured between the manual and Cellpose segmented groups for the same condition (side-by-side orange and blue violin plots).

## Discussion

Autofluorescence microscopy is an attractive tool due to its single cell resolution, sensitivity to cellular function, and nondestructive nature. Due to these advantages, autofluorescence microscopy is a popular method to characterize cell heterogeneity, so it is important to standardize single cell segmentation methods to improve reproducibility and reduce human variability and error. However, cells can be difficult to segment in autofluorescence images where SNR can be low. In such cases, cell borders become less clear, resulting in increased processing times. Fortunately, recent advancements in deep learning have created a host of new tools for single-cell segmentation including Ilastik, Stardist, and Cellpose^8,28–30^. These technologies are easy to train, require minimal adjustments, and promise to automate the task of single-cell segmentation to minimize human error and improve the speed of autofluorescence image analysis.

Commonly, tools for biological image analysis are trained on high contrast images of stained cells, which work well with typical methods of thresholding and edge-detection. In cases of low signal and low SNR like those of autofluorescence which exhibit low quantum yields, these traditional methods are often lacking, require significant amounts of fine tuning, and cannot be used broadly across similar types of images. Cellpose, through its generalist approach, aims to make these models more broadly applicable to a vast array of diverse data sets^8^. While Cellpose is just one algorithm in a crowded landscape of new deep learning tools for image analysis, we can presume that other methods would perform similarly, though Cellpose has made significant effort to ensure that training models is exceedingly simple and accessible^31,32^. Recently, Cellpose has even demonstrated its feasibility to segment three-dimensional objects in an effort to increase its use for additional imaging techniques and analyses, demonstrating continued effort to broaden its application in biomedical research^33,34^. Because Cellpose has already demonstrated the ability to segment a wide range of cell shapes, we attempted to use it for our autofluorescence images and while the base model was adequate, it was clear that improvement was needed. Therefore, we trained our own model.

The capacity of Cellpose to produce comparable masks to human segmentation while significantly reducing the segmentation time demonstrates its utility in automating single-cell segmentation in autofluorescence microscopy. While the Dice Scores may appear low where one would expect a value of 1.0 for a perfect match, a value of 0.76 was previously shown to represent a high level of agreement between two independent human observers^8^. Therefore, in terms of the Dice score, Cellpose appears to perform as well as another human observer **(Fig. 1-2)**. In terms of speed, Cellpose also greatly outperforms human segmentation as expected. While human observers can easily become fatigued segmenting cells and require roughly 11 minutes to segment only 100 regions, Cellpose can segment cells substantially faster **(Supp. Fig. 1)**. Taken together, the speed and reproducibility of Cellpose-generated cell masks greatly reduces the time needed to segment autofluorescence microscopy images.

Importantly, R^2^ values for the autofluorescence FLIM parameters of both 2D and 3D cultures are high, indicating good agreement between manual and Cellpose segmentation for quantitative analysis of autofluorescence FLIM images **(Fig. 3-4)**. While R^2^ values for Cellpose vs manual segmentation were higher for 2D culture compared to 3D PDCOs, this is likely due to the increased presence of irregular cell shapes and blurred boundaries between cells in 3D culture. However, despite the more challenging morphologies associated with 3D organoid culture, Cellpose still retains a similar data distribution as manual masks for each of the three autofluorescence FLIM parameters. While masking of 2D cells was performed across only 9 images, a large selection of images was tested for PDCOs which provided a large sample size for statistically robust results.

While Cellpose provides a high throughput method for single-cell segmentation, it is still important to verify mask accuracy as there is potential for patient-specific differences that may reduce the accuracy of the single-cell masks though we do not anticipate this to be an issue. While our Cellpose segmentation model provides no human-in-the-loop step, some inconsistencies were found in images such as occasional segmentation of the necrotic core and over-segmentation leading to large multi-cell “cells”. Conversely, Cellpose also commonly identifies more nuclei than manual segmentation which could be attributed to human fatigue. Because of the large number of true single nuclei compared to the small numbers of inconsistencies observed, it is likely that these errors are overcome by the large volume of accurate masks. In addition to incorporating a manual check, further refining the Cellpose model may also increase confidence in single-cell segmentation^35^.

Here, we find that decreasing cell segmentation time through a Cellpose model for autofluorescence microscopy can increase the throughput of image analysis with little impact on accuracy. This could enable greater sample sizes in autofluorescence microscopy studies beyond FLIM of NAD(P)H and FAD, including fats like lipofuscin, structural components such as collagen, and commonly occurring pigments like chlorophyl for plant studies. Because FLIM often suffers from low SNR, this study provides confidence not only in the ability of Cellpose to segment microscopy images with high accuracy but also highlights the generalizability of the Cellpose algorithm. Overall, we demonstrated that single-cell distributions of functional variables are highly conserved between manual and Cellpose segmentation methods for autofluorescence microscopy, which provides high confidence in deep learning segmentation methods for low SNR applications such as autofluorescence microscopy.

## Methods

### Cell Culture

#### 2D cell line

PANC-1 cells were obtained from the American Type Culture Collection (ATCC) and cultured in Dulbecco’s minimum essential medium (DMEM) supplemented with 10% Fetal Bovine Serum (FBS) and 1% penicillin-streptomycin (Fisher Scientific). For 2P FLIM, 1.8×10^5^ cells were seeded 48 hours prior to imaging in 35 mm glass-bottom dishes (MatTek). The cells were treated with solutions in Dulbecco’s phosphate-buffered saline (DPBS) of either 4 mM NaCN for 30 mins or 10 mM 2DG for 2 hours prior to imaging. Cells were imaged at three different locations in each dish for an average of 250 cells per treatment group.

#### 3D PDCOs

All PDCO images were acquired from a previously published dataset in Kratz et al.^27^ All studies were completed following Institutional Review Board (IRB) approval with informed consent obtained from subjects through the University of Wisconsin (UW) Molecular Tumor Board Registry (UW IRB#2014-1370) or UW Translational Science BioCore (UW IRB#2016-0934). Cell isolation and organotypic culture techniques were performed as described previously by Kratz et al.^27,36^. Briefly, tissue samples were obtained through endoscopic biopsy or primary surgical resection and treated with chelation buffer. Digestion was performed in DMEM stock with collagenase and dispase. The samples were resuspended in Advanced DMEM/F-12 stock after which organoid suspensions were mixed with Matrigel matrix and plated. Plated cultures on 35 mm glass-bottom dishes (MatTek) were overlaid with 450 μL of feeding medium supplemented with tissue dependent components^36^. PDCO cultures were incubated at 37°C in 5% CO_2_ with medium changes every 48-72 hours and treatment with 10 µmol/L of 5-fluorouracil (5FU), 10 µmol/L 5FU and 40 mmol/L of oxaliplatin (FOLFOX), or control.

### Two-Photon FLIM

#### 2D cell line

Two-photon imaging was performed on a custom-built Multiphoton Imaging System (Bruker) consisting of an inverted microscope (TI-E, Nikon) coupled to an Insight DeepSee+ solid state laser (Spectra Physics). Images were acquired in photon-counting mode using time-correlated single-photon counting electronics (SPC-150, Becker & Hickl GmbH) and GaAsP photomultiplier tubes (H7422P-40, Hamamatsu). Imaging was performed using Prairie View Software (Bruker). The tunable multiphoton laser allowed sequential excitation of NAD(P)H and FAD at 750 and 890 nm, respectively. The emission filter used for NAD(P)H was a bandpass 460/80 nm, whereas FAD used a bandpass 550/100 nm. All samples were imaged using a 40x water immersion, 1.15 NA objective (Plan Apo, Nikon) with an image scan speed of 4.8 μs per pixel, 60 second integration time, and image size of 256 x 256 pixels (2D) or 512×512 pixels (3D). The laser power at the sample was maintained at <10 mW. The instrument response function (IRF) was collected by recording the second harmonic generation signal of urea crystals. A Fluoresbrite YG microsphere (Polysciences Inc.) was imaged as a daily standard for fluorescence lifetime. The lifetime decay curves for these beads were fit to a single exponential decay, and the fluorescence lifetime was measured to be 2.1 ns (n = 7), which is consistent with published values^37,38^.

#### 3D PDCOs

All PDCO images were acquired from a previously published dataset in Kratz et al.^27^ Briefly, imaging protocol was the same as for PANC-1, though differences existed for laser [Mai Tai DeepSee Ti:sapphire laser (Coherent, Inc)] and NAD(P)H bandpass filter (440/80 nm).

### Cellpose Model Development

Cellpose is a versatile algorithm for cell and nucleus segmentation, featuring pre-trained models developed on a wide range of microscopy images. It also supports a human-in-the-loop pipeline for the rapid prototyping of new custom models^31^. However, the pre-trained models were unable to accurately segment the nuclei and cells in our autofluorescence microscopy images.

To train a new deep learning model in Cellpose, annotated data is required. Labeled images where each object has a unique label (masks) paired with the corresponding autofluorescence image were used for this purpose. To streamline the annotation process, we cropped, without rescaling, segments from at least three different images from 3D cell cultures, each containing 5-10 cells. This approach was intended to save time during annotation. For each cell in these cropped images, we drew the nucleus masks using the graphical user interface (GUI) of Cellpose.

These pairs of images and masks (cropped image with nucleus mask) were used to train a new model for segmentation of nuclei from 3D cell culture images. Following the methodology described in the “Cellpose 2.0: how to train your own model” paper^31^. The training parameters were as follows: --pretrained_model cyto, --chan 2, --chan2 1, --learning_rate 0.01, -- weight_decay 0.0001, and --n_epochs 500.

### Image analysis

Fluorescence lifetimes were extracted using SPCImage software (SPCImage v8.1, Becker & Hickl). The fluorescence lifetime decay curve was convolved with the IRF, using an iterative parameter optimization to obtain the lowest sum of the squared differences between model and data (weighted least squares algorithm), the goodness of fit was monitored to ensure a chisquared value < 1.3. The two-component exponential decay model is *I(t) = α*_*1*_^*-t/τ*^_*2*_ + *α*_*2*_^*-t/τ*^_*2*_ + *C*, where *I(t)* is the fluorescence intensity at time *t, α* is the fractional contribution of each component, *τ* is the lifetime of each component, and *C* accounts for background signal. A bin of 3×3 pixels was used to increase photon counts for decays. The two lifetime components are used to distinguish between the free and bound forms of NAD(P)H and FAD^9,10^. The mean fluorescence lifetime was calculated using *τ*_*m*_ *= α*_*1*_*τ*_*1*_ + *α*_*2*_*τ*_*2*_. Additionally, the decay curves for NAD(P)H and FAD were integrated for each pixel to obtain intensity values. These intensity values are used to calculate the ORR using the equation *ORR = I*_*NAD(P)H*_ */ [I*_*NAD(P)H*_ + *I*_*FAD*_], where the intensities of NAD(P)H and FAD are given by *I*_*NAD(P)H*_ and *I*_*FAD*_, respectively.

### Image segmentation

Manual segmentation was performed by human observers who identified and manually circled each nucleus within each image using CellProfiler software. The individuals tasked with performing image segmentation underwent a comprehensive training regimen designed to enhance accuracy and precision in segmenting relevant features within NAD(P)H intensity images. The training encompassed observing a series of annotated NAD(P)H intensity images spanning 2D and 3D cultures and then segmenting those images with guidance given through comparisons of the annotated images with their segmentation. After manual identification of nuclei, the cytoplasm region was isolated using a Python script to propagate out from the masked nuclei by 8 µm (10 pixels) to match the protocol of prior studies where we employed manual segmentation^23,27^. The reported segmentation time was determined using a stopwatch from the opening of an image set to the closing of an image set and dividing by the number of images in the set.

Cellpose segmentation was performed by loading images into Cellpose where the software identified nuclear regions using the model described above. No further corrections were made to the masks. The cytoplasm region was then recognized using a Python script to propagate out from the nuclei by 8 µm (10 pixels) to match the procedure of the manual segmentation above. Segmentation time was determined using the Python DateTime module to measure the execution time of the segmentation script^39^.

For additional validation, after initial segmentation, all PANC-1 images were rotated 90 degrees and segmented again by the same observer. These rotated images were presented as new images to the observer, who were unaware that they were re-segmenting the same images. Patient 6 PDCO images were also rotated and manually segmented again, though not by the same observer.

Finally, Cell-analysis-tools 0.0.14 was used for feature extraction of NAD(P)H and FAD autofluorescence imaging features (*τ*_*m*_, *τ*_*1*_, *τ*_*2*_, *α*_*1*_, *α*_*2*_, *I, ORR*) which were averaged across all pixels within the cytoplasm mask of each segmented cell^40^. This method was used for both the manual and Cellpose segmented images.

### Statistics

Differences between Dice scores were tested using an Ordinary one-way ANOVA test. Effect sizes between groups in NAD(P)H τ_m_, FAD τ_m_, and ORR were calculated with Glass’s Delta because comparisons of very large sample sizes of individual cells always pass traditional significance tests unless the population effect size is truly zero^41^. Glass’s Delta is defined as *Δ = (µ*_*control*_ *-µ*_*test*_*) / σ*_*control*_ *where* values ranging from 0-0.2 represent no change, 0.2-0.5 small change, 0.5-0.8 represent moderate change, and anything higher than 0.8 signifies a large change^42,43^. Simple linear regression modeling was performed using GraphPad Prism 9.4.1.

## Acknowledgements

We thank Kratz et al.^27^ for providing the organoid FLIM images as well as their manually segmented nuclear masks. Additionally, we would like to thank the current members of the Skala and Deming labs for the discussions that led to the improvements in our computational methods and the refinement of our analytical techniques. We also thank graphic designer, Matthew Stefely, for assistance in finalizing our figures.

## Author Contributions

JMR: design, analysis, interpretation, drafted the work; AAG: conception, design, interpretation; ECG: analysis; MCC: creation of new software used in the work; AK: acquisition, analysis; SU and AS: interpretation; DD: conception, design, interpretation of data; BC: design, interpretation of data; MCS: conception, design, interpretation of data, drafted the work. All authors have read and approved the final manuscript.

## Code, Data, and Materials Availability

The repository with the initial models and pipelines is located here: https://github.com/COBA-NIH/SkalaLab

## Funding

P41 GM135019, R37 CA226526, R01 CA272855, P30 CA014520 (Core Grant, University of Wisconsin Carbone Cancer Center), ACI/Schwenn Family Professorship, JD Fluno Family Colorectal Cancer Precision Medicine Program, Carol Skornicka Chair, Morgridge Institute for Research.

## Notes

https://github.com/COBA-NIH/SkalaLab

### Competing Interest Statement

The authors have declared no competing interest.

