## Supplementary material for "Cellpose as a reliable method for single-cell segmentation of autofluorescence microscopy images": Supp. Fig. 1

### Slide 1
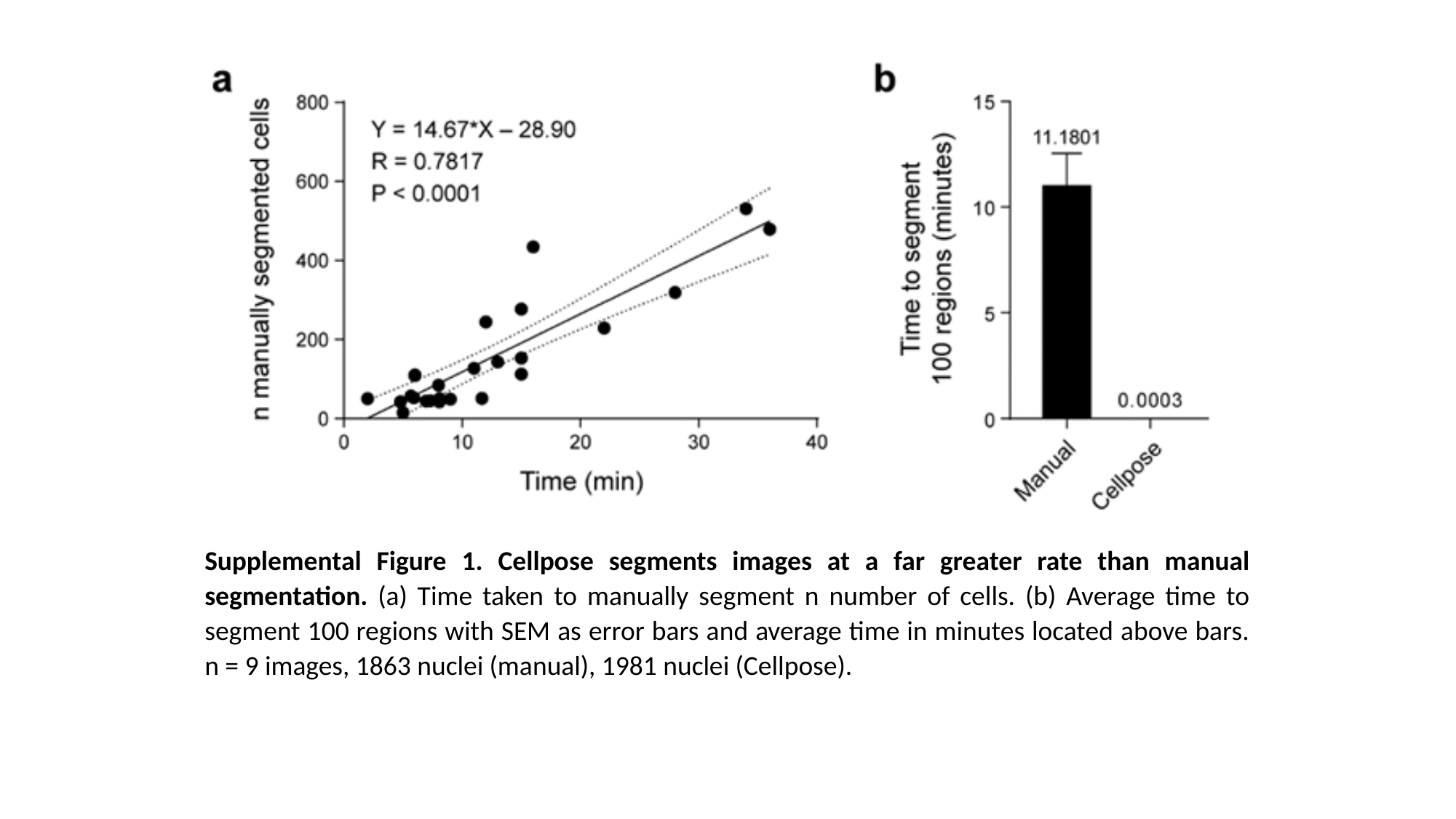

Supplemental Figure 1. Cellpose segments images at a far greater rate than manual segmentation. (a) Time taken to manually segment n number of cells. (b) Average time to segment 100 regions with SEM as error bars and average time in minutes located above bars. n = 9 images, 1863 nuclei (manual), 1981 nuclei (Cellpose).
