## Supplementary material for "Cellpose as a reliable method for single-cell segmentation of autofluorescence microscopy images": Supp. Fig. 2

### Slide 1
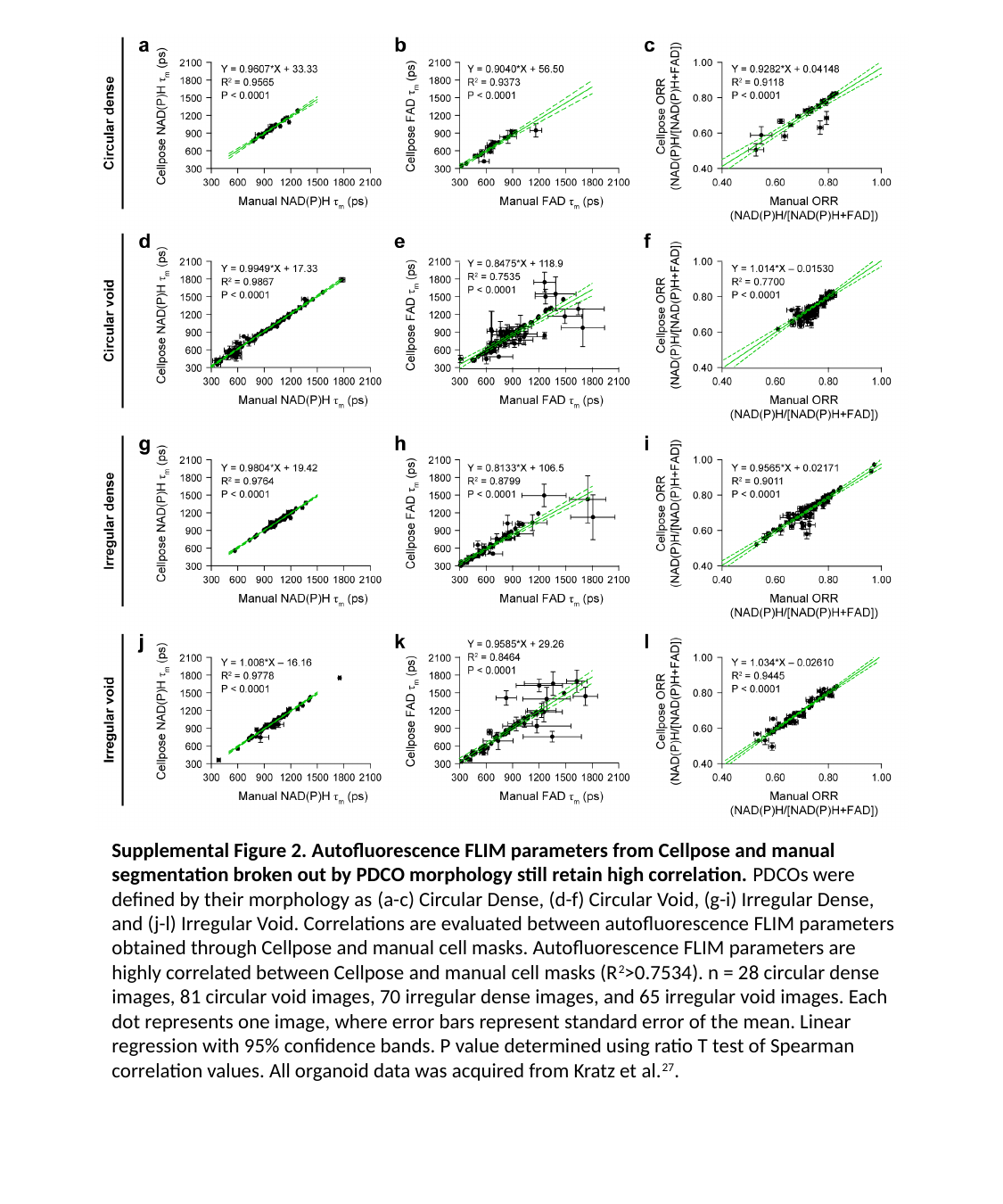

Supplemental Figure 2. Autofluorescence FLIM parameters from Cellpose and manual segmentation broken out by PDCO morphology still retain high correlation. PDCOs were defined by their morphology as (a-c) Circular Dense, (d-f) Circular Void, (g-i) Irregular Dense, and (j-l) Irregular Void. Correlations are evaluated between autofluorescence FLIM parameters obtained through Cellpose and manual cell masks. Autofluorescence FLIM parameters are highly correlated between Cellpose and manual cell masks (R^2^>0.7534). n = 28 circular dense images, 81 circular void images, 70 irregular dense images, and 65 irregular void images. Each dot represents one image, where error bars represent standard error of the mean. Linear regression with 95% confidence bands. P value determined using ratio T test of Spearman correlation values. All organoid data was acquired from Kratz et al.^27^.
