## Supplementary material for "Cellpose as a reliable method for single-cell segmentation of autofluorescence microscopy images": Supp. Fig. 3

### Slide 1
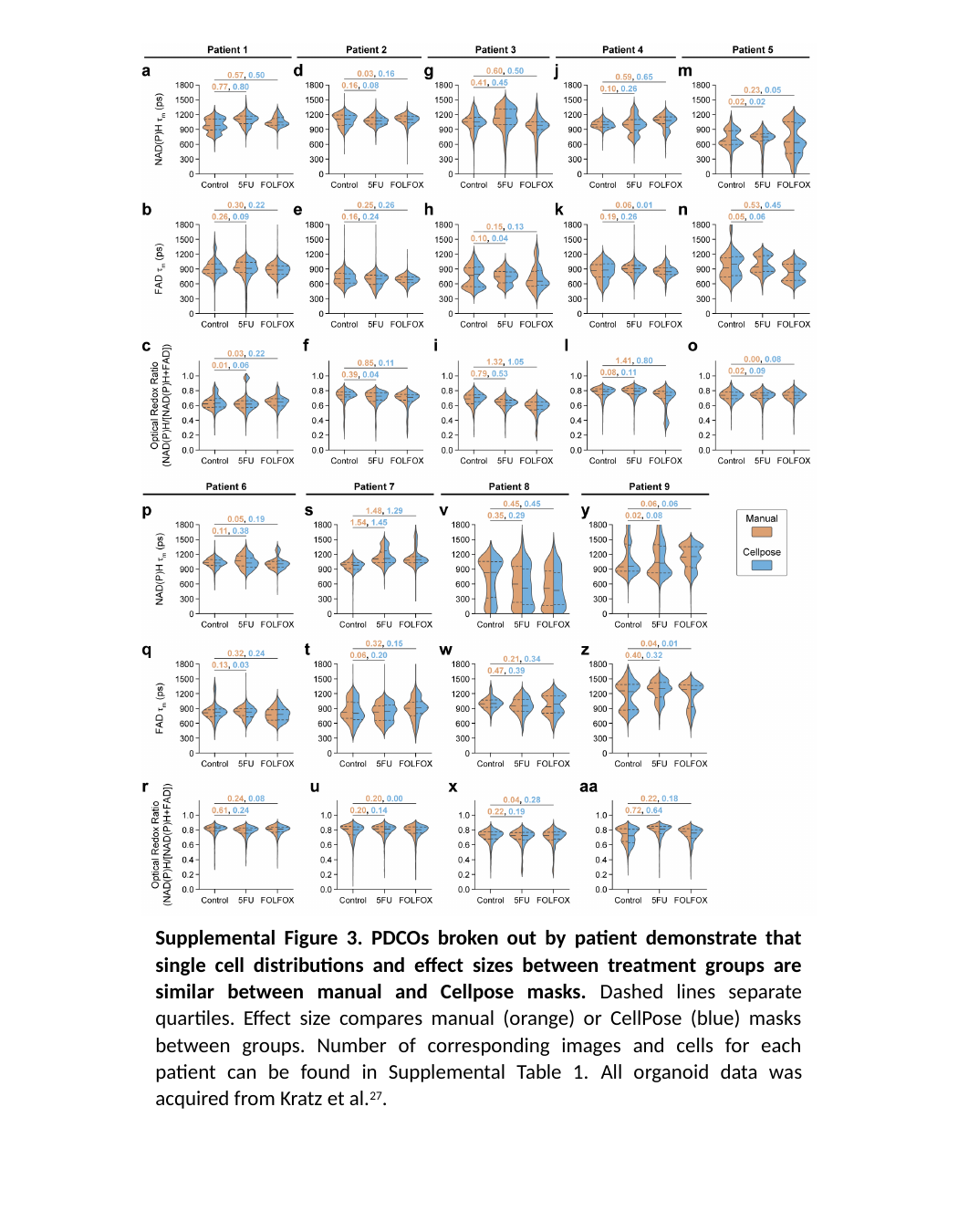

Supplemental Figure 3. PDCOs broken out by patient demonstrate that single cell distributions and effect sizes between treatment groups are similar between manual and Cellpose masks. Dashed lines separate quartiles. Effect size compares manual (orange) or CellPose (blue) masks between groups. Number of corresponding images and cells for each patient can be found in Supplemental Table 1. All organoid data was acquired from Kratz et al.^27^.
