## Supplementary material for "Cellpose as a reliable method for single-cell segmentation of autofluorescence microscopy images": Supp. Table 1

### Slide 1
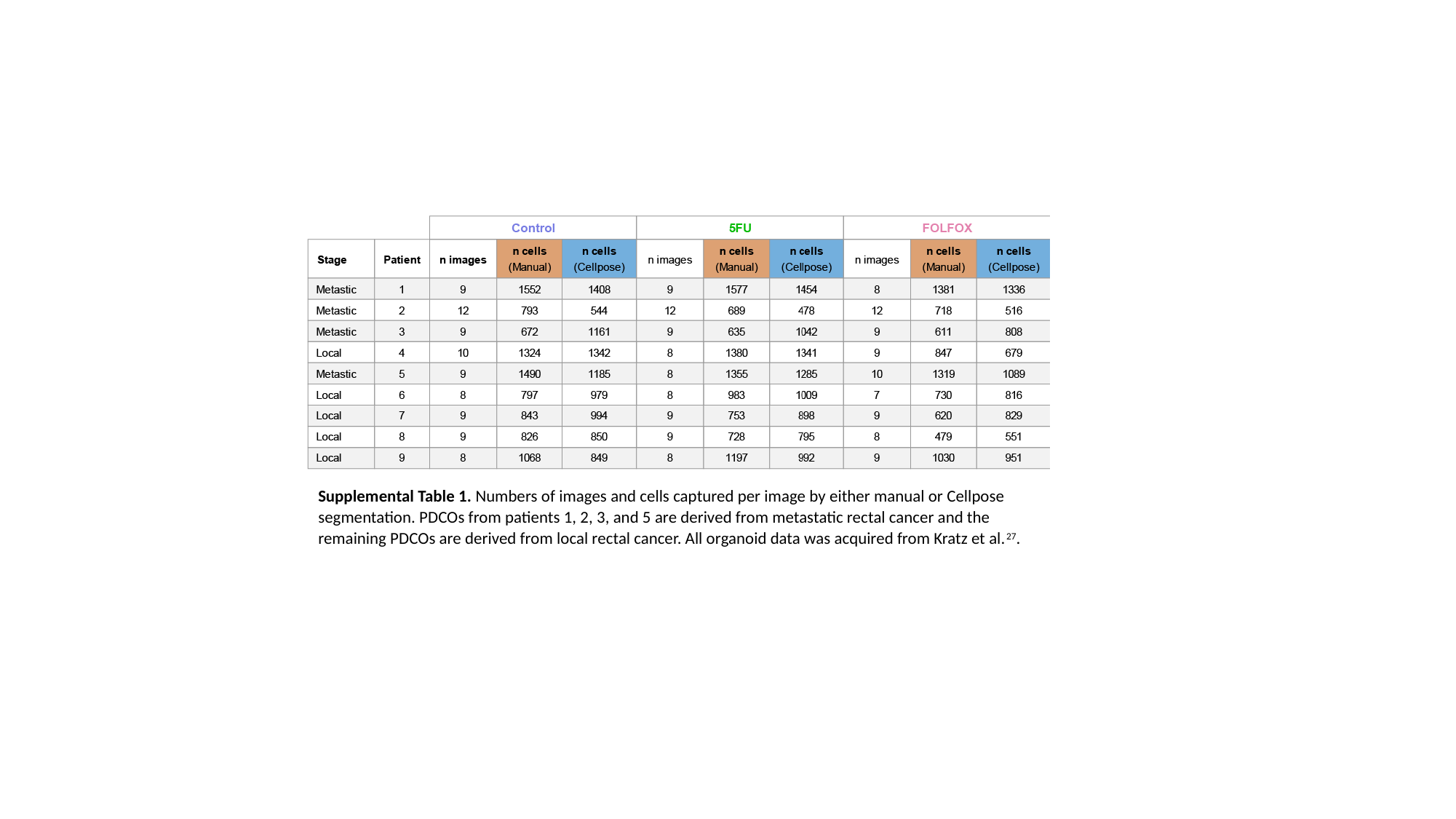

Supplemental Table 1. Numbers of images and cells captured per image by either manual or Cellpose segmentation. PDCOs from patients 1, 2, 3, and 5 are derived from metastatic rectal cancer and the remaining PDCOs are derived from local rectal cancer. All organoid data was acquired from Kratz et al.^27^.
